# Decoding Chronic Pain States from Distributed Intracranial Recordings

**DOI:** 10.64898/2026.06.16.732555

**Authors:** Jeremy Saal, Ankit N. Khambhati, Edward F. Chang, Prasad Shirvalkar

**Affiliations:** UCSF Weill Institute for Neurosciences, University of California San Francisco, San Francisco, CA, USA; UCSF Department of Anesthesiology and Perioperative Care, Division of Pain Medicine, University of California San Francisco, San Francisco, CA, USA; UCSF Department of Neurology, University of California San Francisco, San Francisco, CA, USA; UCSF Department of Neurological Surgery, University of California San Francisco, San Francisco, CA, USA; UCSF Department of Physiology, University of California San Francisco, San Francisco, CA, USA

## Abstract

Chronic pain engages distributed cortical and subcortical circuits, and large-scale intracranial recordings in humans offer a valuable opportunity to characterize its neural signatures. Here, we recorded multi-day stereoelectroencephalography (sEEG) from six participants with refractory chronic neuropathic pain, each implanted with sEEG electrodes spanning dozens of cortical and subcortical structures. Using simultaneous chronic pain ratings, we decoded spontaneous high versus low pain states within individuals (median area under the curve = 0.72; five of six participants performed above chance). Pain-predictive signals were broadly distributed and highly participant-specific. However, mapping the spatial distribution of pain-predictive features revealed preferential representation within canonical macroscale networks: beta-band activity in the default mode network and high-gamma activity in the salience network. These results demonstrate that intracranial recordings can capture distributed, network-organized representations of spontaneous chronic pain states.

## Introduction

Chronic pain is a leading cause of disability worldwide (1). While recent work shows that chronic pain states can be decoded from intracranial neural activity (2–4), it remains unclear which regions and oscillatory features carry pain-state information in intracranial recordings, and how this information is organized across the brain. Neuroimaging implicates a widely distributed set of cortical and subcortical structures (5), but whether pain representation is widely distributed at the level of local field potentials is unknown. Non-invasive imaging has implicated the default mode and salience networks in chronic pain (6,7), but whether these network-level signatures fluctuate with pain state and emerge in intracranial recordings remains an open question. Finally, both neural signals and pain intensity fluctuate slowly over multi-day recordings, producing both short-term autocorrelations and long-term trends whose relevance to decoding performance is poorly understood. Resolving these questions is necessary to move beyond proof-of-concept decoding and characterize how pain is represented in intracranial recordings.

Multivariate fMRI analyses have been used to predict acute experimental pain intensity from distributed activation patterns with high sensitivity (8), and subsequent work has identified functional connectivity signatures that generalize to tonic and clinical pain (9). Personalized fMRI models can track spontaneous pain fluctuations within individuals, but do not generalize across patients, suggesting that the neural signatures of chronic pain are individually unique (10). However, fMRI is limited by measuring hemodynamic changes rather than direct neural activity, which may obscure the underlying neural signals. Direct intracranial recordings address this limitation by providing access to local field potentials (LFPs), which directly reflect local neural activity. Early studies employing intracranial methods identified pain-related spectral correlates in the sensory thalamus and periaqueductal gray but did not test whether pain state could be predicted from neural activity(11–13). More recently, intracranial work has established the feasibility of decoding acute and chronic pain from distributed recordings (2–4,14). However, these studies leave the questions outlined above largely unresolved.

In this study, we characterize the representation of spontaneous chronic pain states across the brain in human participants. We analyzed stereoelectroencephalography (sEEG) recordings from six participants with refractory chronic neuropathic pain, including post-stroke pain, complex regional pain syndrome, chemotherapy-induced neuropathy, and spinal cord injury. Each participant was implanted with sEEG electrodes spanning dozens of cortical and subcortical structures, including the anterior cingulate cortex, insula, thalamus, dorsomedial prefrontal cortex (DMPFC), dorsolateral prefrontal cortex (DLPFC), somatosensory cortex, and basal ganglia. Leveraging multivariate and univariate decoding approaches and a Leave-2-Days-Out cross-validation scheme that prevents within-day temporal leakage, we map how pain is represented across brain regions, spectral frequency bands, and canonical macroscale networks.

Multivariate decoding of chronic pain states was robust across multi-day timescales under cross-validation that accounted for autocorrelation. Electrode-contact-level univariate models revealed broadly distributed pain-predictive features, with beta and gamma bands yielding the strongest decoding performance and the most informative brain regions varying across individuals. Despite this variability, the right DMPFC was the most consistent pain-predictive region amongst participants with significant decoding, a convergence enabled by its consistent coverage across the cohort. Mapping decoding performance onto the Yeo 7-network parcellation revealed that pain-predictive beta activity was enriched in the default mode network, while pain-predictive high-gamma activity was enriched in the salience network (15). Together, these results demonstrate that pain-predictive regions are broadly distributed and participant-specific, while cortical signals are preferentially enriched within the default mode and salience networks.

## Results

### Participant overview and pain characterization

We analyzed intracranial recordings from six participants with chronic neuropathic pain (P1–P6) undergoing inpatient sEEG mapping as part of a closed-loop deep brain stimulation (DBS) trial (see Methods). Each participant was implanted with 8–11 sEEG electrodes targeting both cortical and subcortical structures, such as the anterior cingulate cortex, insula, thalamus, prefrontal cortex, somatosensory cortex, and basal ganglia (Fig. 1a, Supplementary Fig. S1). Four participants also had electrocorticography (ECoG) strips covering the motor and sensory cortices (P1–P4). Only contacts within gray matter were included in this study. Participants then underwent 8–10 days of continuous neural recording in the hospital. During this period, we tested deep brain stimulation parameters and collected pain surveys. Participants completed 57–105 pain surveys (Fig. 1b) using a pain intensity visual analog scale (VAS, 0–100, Fig. 1c).

**Figure 1.**
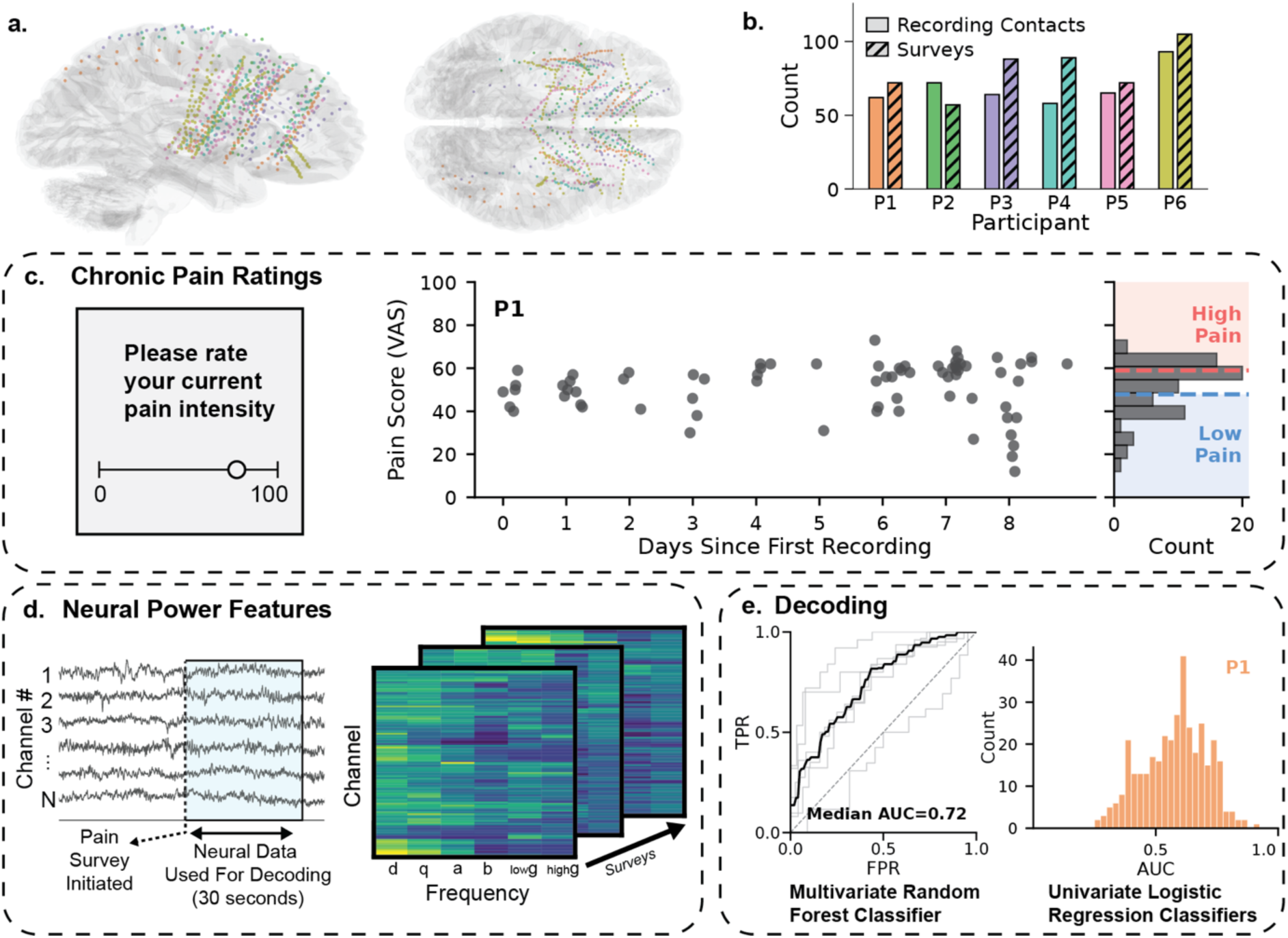
Study design and decoding pipeline. (a) Glass brain rendering showing stereoelectroencephalography (sEEG) contact coverage, colored by participant (colors as per b). (b) Number of sEEG recording contacts and surveys per participant. Solid and hatched bars show the number of sEEG recording contacts and the number of surveys collected during each recording period, respectively. (c) Chronic pain ratings. Left, schematic showing the visual analog scale (VAS) participants used to rate their pain intensity. Right, scatter plot showing VAS pain intensity scores of example participant P1 over approximately nine days. Marginal histogram shows the distribution of pain scores and the binarization thresholds (33rd and 67th percentiles) used to determine high and low pain labels for decoding. (d) Neural power features used for decoding. Left, schematic of sEEG signals showing the 30-second window of neural data extracted after each pain survey was initiated. Right, heatmap showing spectral power features extracted per survey: delta, theta, alpha, beta, low gamma, and high gamma power were extracted per contact. (e) Decoding outputs. Left, ROC curves from the multivariate random forest classifier (thin gray lines, individual participants; bold black line, pointwise median TPR across participants; median AUC = 0.72). Right, distribution of cross-validated AUCs from example participant P1’s univariate logistic regression classifiers (one per contact-frequency combination).

Median pain intensity scores across participants ranged from 15 (P5) to 82 (P6) on the VAS (Supplementary Fig. S2, Supplementary Table S1). Participants gave pain reports with a median inter-survey interval of 0.3 to 1.2 hours. We dichotomized pain scores within each participant, with the bottom 33% of scores considered low pain and the top 33% high pain. We chose binary classification to reflect the threshold-crossing paradigm required by onboard closed-loop DBS devices. This percentile-based grouping preserved class balance, yielding 22–40 trials per class per participant. Participants had between 57 and 90 gray matter contacts each (Fig. 1b).

### Chronic pain states are robustly decodable from multi-site sEEG

We first asked whether low versus high pain states can be decoded using power features from sEEG signals during each survey (Fig. 1d), employing a cross-validation scheme designed to prevent within-day temporal leakage due to temporal autocorrelation of both pain ratings and neural activity. A key challenge in multi-day inpatient recordings is that both neural signals and pain intensity fluctuate slowly over hours to days, driven by factors such as medication timing and the natural time course of chronic pain. If temporally adjacent samples appear in both training and test sets, a classifier can exploit this shared temporal structure rather than learning genuine pain-related features, producing inflated performance estimates. To account for this, we employed a Leave-2-Days-Out cross-validation scheme, in which all samples from two held-out days formed the test set while the remaining days were used for training, ensuring that no temporal adjacency existed between training and test observations (Supplementary Fig. S3; see Methods). Surveys occurring within 5 minutes of stimulation were also excluded to minimize carryover effects. We used a random forest classifier on region-level spectral features derived from principal component analysis (PCA) by retaining the top principal component per region as predictor variables (see Methods, Fig. 1e, 2a). Across participants, we observed a median area under the receiver operating characteristic curve (AUC) of 0.72 (mean 0.70, range 0.41–0.91) (Fig. 2b). We used permutation testing (1,000 label shuffles) to assess significance. We found that five out of six participants achieved above-chance performance (false discovery rate (FDR)-corrected p < 0.05; Fig. 2c; predictions for an example participant shown in Fig. 2d; all participants in Supplementary Fig. S4). P2 was the only participant for whom decoding did not surpass chance level (AUC = 0.41, p = 0.79), likely attributable to the predominantly peripheral mechanisms of their chemotherapy-induced peripheral neuropathy (see Discussion). Across the five significant participants, decoding sensitivity ranged from 0.70 (P3) to 0.92 (P5) and specificity from 0.57 (P4) to 0.93 (P3) (Fig. 2e).

**Figure 2.**
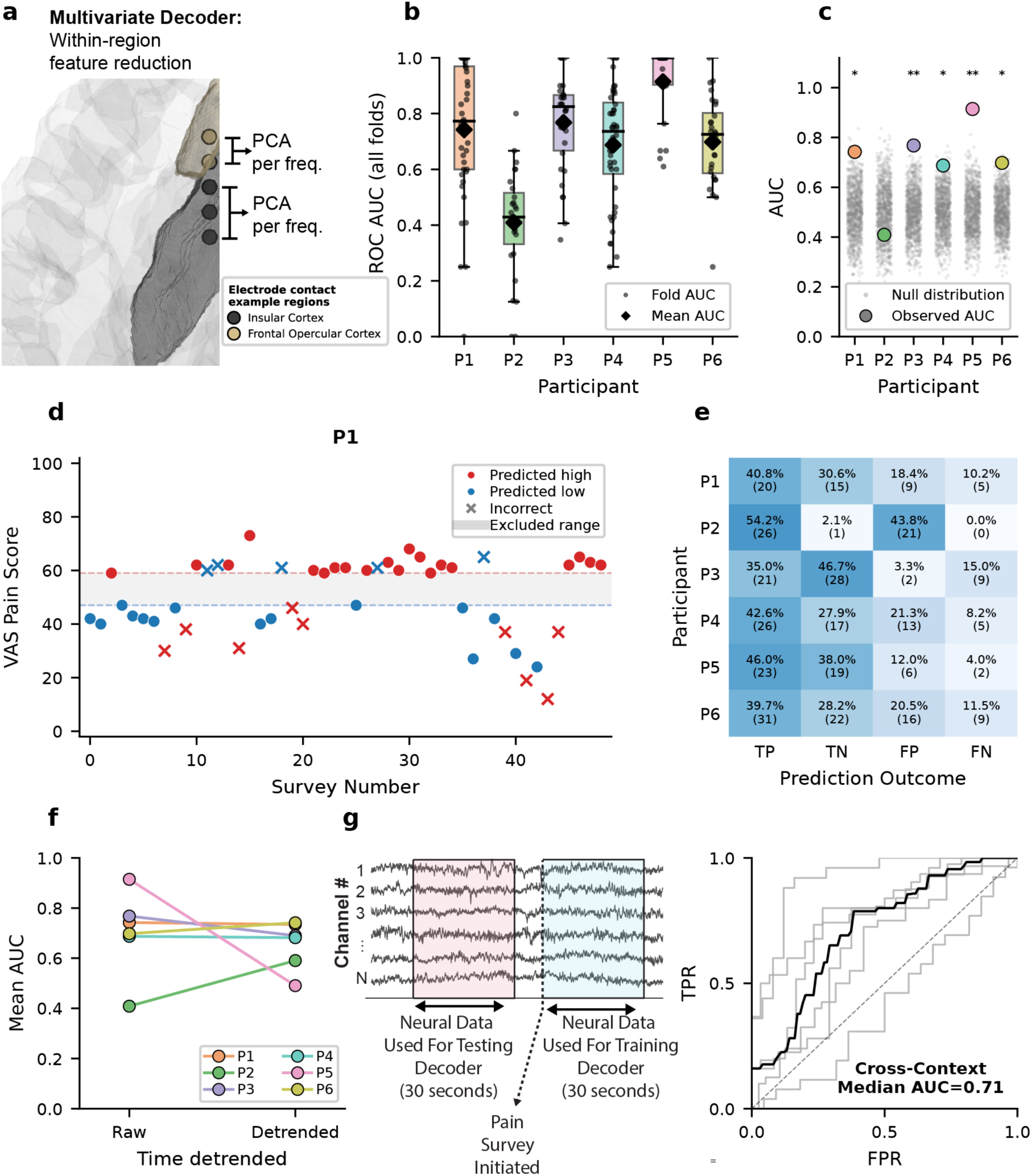
Multivariate decoding of chronic pain states. (a) Multivariate decoder feature reduction. Schematic illustrating region-level feature extraction, with the insular cortex and frontal opercular cortex shown as example regions. Within each anatomical region and frequency band, contact-level spectral power features are reduced to a single region-level feature via principal component analysis (PCA). (b) Cross-validated ROC AUC values from each Leave-2-Days-Out fold (n = 6 participants). Box edges, 25th–75th percentiles; center line, median; whiskers extend to the most extreme value within 1.5x IQR. Small dots, individual fold AUCs; black diamonds, per-participant mean AUC. (c) Permutation significance testing for the multivariate random forest decoder. Gray points, null AUC distribution from 1,000 label-shuffled permutations per participant; colored points, observed AUC. Asterisks denote significance after Benjamini-Hochberg FDR correction (*p < 0.05, **p < 0.01, ***p < 0.001). Five of six participants showed above-chance-level decoding. (d) Example per-survey predictions for P1. The middle third of VAS scores was excluded during binarization. Filled circles, correct predictions; crosses, incorrect predictions; red points, high-pain predictions; blue points, low-pain predictions. (e) Per-participant confusion values showing the proportion of true positives, true negatives, false positives, and false negatives. (f) Decoding performance before and after removing linear temporal trends. Left, raw data; right, after detrending. Each point is one participant’s mean AUC across Leave-2-Days-Out folds. After detrending, decoding remained significant in four of the five initially significant participants; P5 no longer exceeded chance, and P2 remained non-significant. (g) Cross-context validation. Left, schematic showing that neural data extracted while participants completed surveys was used to train the classifier and tested on neural data from the 30-second window beginning one minute before each survey. Right, cross-context ROC curves: thin gray lines, individual participants (n = 6); bold black line, pointwise median TPR across participants (interpolated to a common FPR grid). Median AUC = 0.71, comparable to within-context decoding (median 0.72), indicating that decoding performance is not specific to survey-taking behavior.

We performed multiple control analyses to ensure robustness against potential temporal and contextual confounds. If efficacious stimulation sites are discovered early and continuously applied, pain scores may decrease progressively over the recording period, as was observed in P5 (Supplementary Fig. S2). Furthermore, electrode impedance changes over time can introduce nonspecific drift in neural features. These co-occurring trends could give rise to an apparent relationship between neural activity and pain that does not reflect genuine pain-related neural activity. To address this, we repeated the multivariate analysis after removing linear temporal trends from the spectral features. Among the five initially significant participants, performance remained above chance after detrending in all except P5; P2 remained non-significant (permutation test, p > 0.05; Fig. 2f). These results indicate that temporal drift was correlated with pain intensity in this participant but did not account for the overall pattern of significant decoding across the cohort. Finally, we asked whether pain-related neural signatures are tonic representations that persist beyond the survey period, or whether they are evoked by the act of introspection during survey-taking. To test this, we used a cross-context approach in which we trained our decoder on during-survey data but tested it on neural signals from a 30-second window beginning one minute prior to the survey (Fig. 2g, left). Cross-context performance was comparable to within-context performance (median AUC of 0.71 in cross-context vs. 0.72 in baseline; Fig. 2g, right), suggesting that pain-related neural signatures reflect ongoing brain states rather than transient, task-evoked activity tied to the survey-taking process. Together, these analyses indicate that the main decoding result is not readily explained by temporal leakage, acute stimulation carryover, or survey-context effects in most participants.

While multivariate models performed well, multicollinearity among features limits the interpretability of feature importances, as correlated features can substitute for one another, even with regularization. Median pairwise correlations between features ranged from 0.20 (P4) to 0.36 (P3) (Supplementary Fig. S5), and only 20–35 effective dimensions out of the 120–204 features were needed (15–18% of the total feature count; see Methods). Consistent with this redundancy, permutation importance was low even for the top-ranked features in each participant (Supplementary Fig. S6). Moreover, noisy neural features can appear important solely because they reduce noise in other features rather than because they independently carry pain-relevant information. For clinical applications, such as closed-loop DBS target selection, this distinction is critical: a region flagged as important by a multivariate model may not itself carry pain-relevant information. Together, these multivariate results establish that chronic pain states are reliably represented in multi-site sEEG recordings, and these representations are stable across multiple days. However, the observed multicollinearity among region-level features limits interpretability of feature importance, motivating the complementary univariate analyses that follow.

### Pain-informative signals are broadly distributed across cortical and subcortical sites

We next asked where pain-informative signals are localized across the brain. We expected anatomical regions providing strong decoding performance to be largely confined to the descending pain modulatory pathway (e.g., the anterior cingulate cortex, prefrontal cortex, and periaqueductal gray) but found pain-informative signals to be more broadly distributed. We trained univariate logistic regression per gray matter electrode contact and frequency band (Fig. 3a), using Leave-2-Days-Out cross-validation (Supplementary Fig. S3). This yielded a total model count ranging from 342 to 540 across participants. Although the median AUC across univariate models was modest (range 0.48–0.67 across participants), a substantial proportion of univariate models showed evidence of pain-informative signals (high AUC) per participant (Fig. 3b). To identify specific regions with statistically reliable pain-informative signals, we characterized performance at the contact level; a contact was considered significant if at least one of its six frequency-band models survived Benjamini–Hochberg FDR correction across those six tests (Fig. 3c, d). Applying this correction, the per-participant proportion of contacts with significant decoding in at least one frequency band (excluding P2 due to insignificant multivariate decoding) ranged from 38% (P6) to 87% (P3), indicating that pain-related spectral features were broadly distributed rather than confined to a single anatomical locus. Specifically, 28/60 contacts (47%) were significant in P1, 0/68 (0%) in P2, 53/61 (87%) in P3, 25/57 (44%) in P4, 55/64 (86%) in P5, and 34/90 (38%) in P6. Together, these results indicate that pain-informative oscillatory signals are broadly distributed across cortical and subcortical contacts rather than confined to a single anatomical locus.

**Figure 3.**
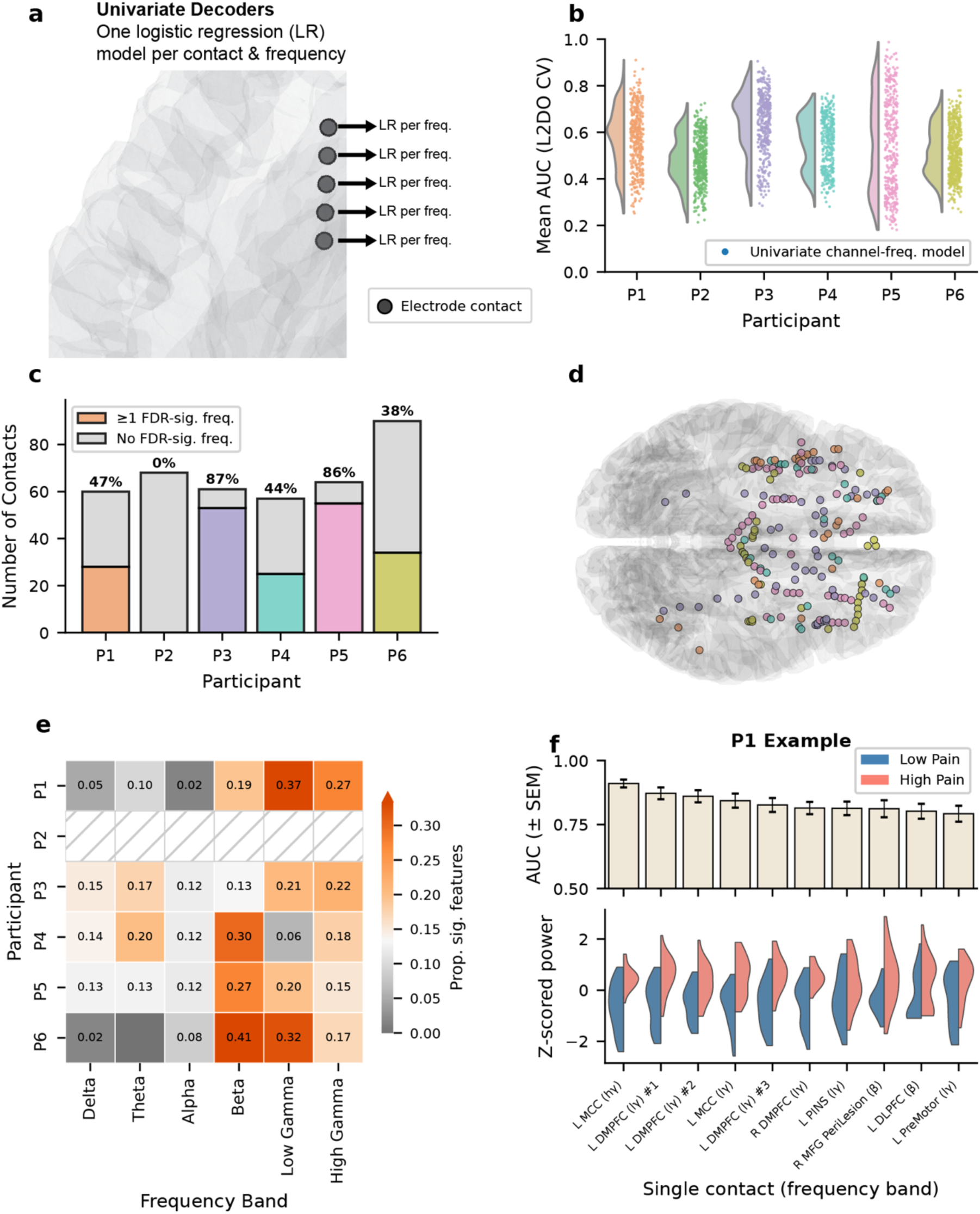
Univariate decoding reveals distributed pain-relevant neural activity. (a) Univariate decoder schematic. For each electrode contact and frequency band, a separate logistic regression (LR) classifier was trained, yielding one model per contact–frequency combination. (b) Distributions of cross-validated AUCs across all contact–frequency models for each participant (raincloud plots: half-violin density with individual model AUCs as dots). (c) Per-participant proportion of contacts with at least one FDR-significant frequency band (Benjamini-Hochberg correction across the six bands tested per contact). (d) Contacts surviving FDR correction in (c), rendered on a 3D glass brain and colored by participant. (e) Heatmap showing the within-participant proportion of FDR-significant univariate features in each frequency band. Colormap is centered on the proportion expected from a uniform distribution across bands. Rows sum to one. Hatched row indicates P2, who did not have any significant decoding models. (f) Example participant (P1). Top, mean AUC ± SEM across Leave-2-Days-Out folds for the top-ranked single-contact contact-frequency features (sorted in descending order). Bottom, split-violin distributions of z-scored spectral power for low-pain (blue) and high-pain (red) observations at each corresponding contact.

To assess whether pain-predictive features were preferentially represented in certain frequency bands, we examined the distribution of each participant’s FDR-significant univariate features across the six bands tested and computed the within-participant proportion of significant features in each band. Across cortical and subcortical contacts, higher-frequency bands (beta, low gamma, and high gamma) consistently carried the strongest univariate decoding signal across participants (Fig. 3e). The dominant band was participant-specific: beta in P4, P5, and P6; low gamma in P1 (Fig. 3f); high gamma in P3; P2 had no FDR-significant features.

The highest-performing univariate contact-based models spanned multiple anatomical regions and were participant-specific (Fig. 4, Supplementary Fig. S7). In P1, top contacts were concentrated in the left midcingulate cortex (MCC) and left dorsomedial prefrontal cortex (DMPFC), predominantly in the low gamma and high-gamma bands (Fig. 3f). In P3, the right frontal operculum (FOP), right DMPFC, and right anterior insula (AINS) contributed the strongest decoding, with multiple insular contacts (exceeding AUC 0.85 in the gamma bands). In P4, top contacts were distributed across the DMPFC, thalamus, and dorsolateral prefrontal cortex (DLPFC), with theta, alpha, and beta bands predominating. In P5, subcortical structures drove the strongest decoding: the best single univariate model was in the right putamen (high gamma, AUC = 0.99), followed by contacts in the DLPFC, thalamus, and periventricular gray (PVG). However, P5 decoding did not survive detrending (Fig. 2f), so these anatomical results should be interpreted with caution. In P6, the top contacts were in the PVG and orbitofrontal cortex (OFC). Together, these results show the individualized nature of pain-predictive features across our participants.

**Figure 4.**
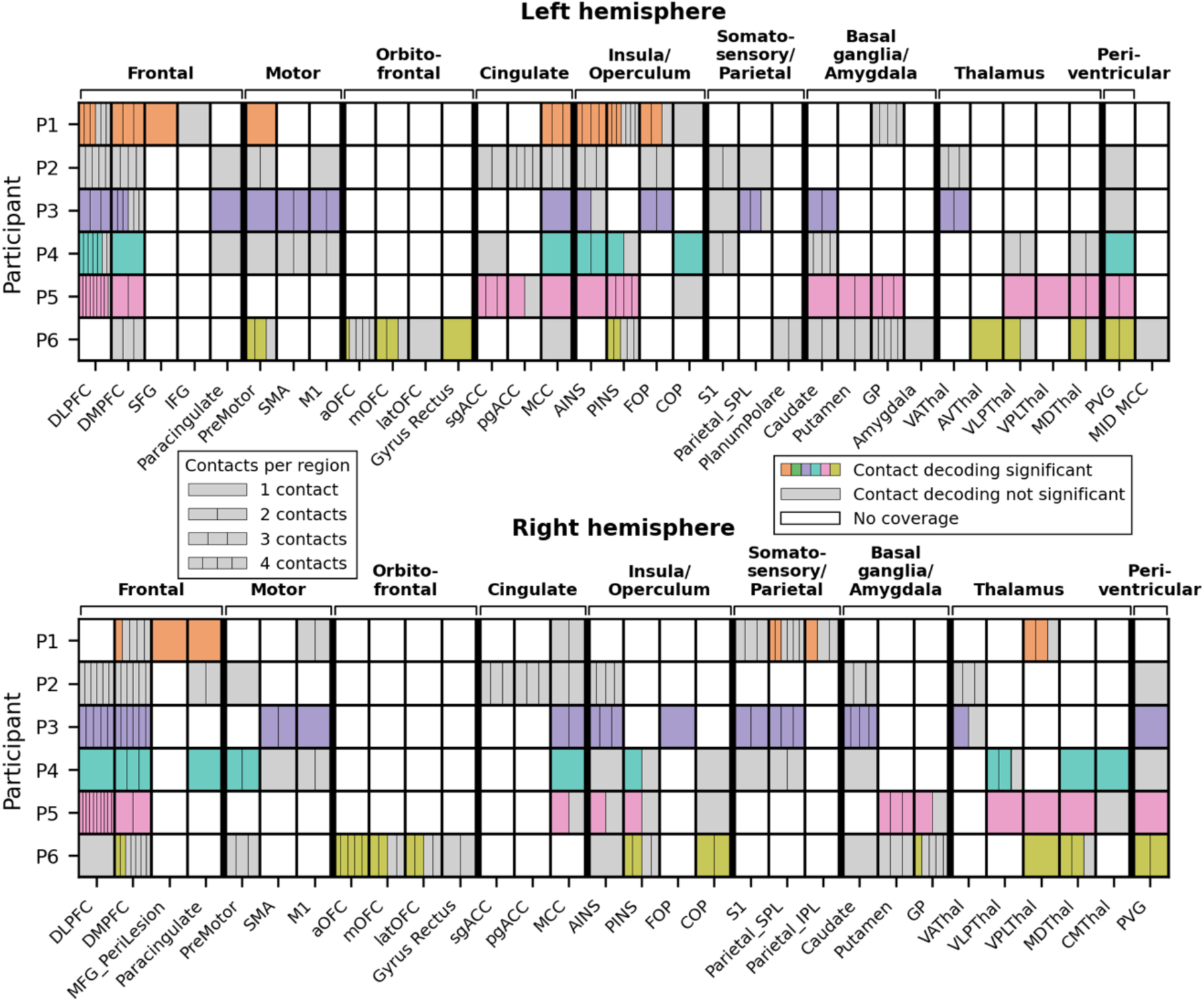
Anatomical distribution of pain-informative contacts across participants. One grid is displayed per hemisphere: left hemisphere (top) and right hemisphere (bottom). Each grid has rows for participants P1–P6 and columns for anatomical regions grouped by coarse anatomical bracket. Within each cell, individual contacts are shown as sub-tiles (legend displays up to four; some regions contain more). Contacts with at least one frequency band surviving Benjamini-Hochberg FDR correction are colored by participant. Non-significant contacts are shown in gray, and regions without coverage are shown in white. Anatomical regions abbreviated as: AINS, anterior insula; aOFC, anterior orbitofrontal cortex; AVThal, anterior ventral thalamus; CMThal, centromedian thalamus; COP, central operculum; DLPFC, dorsolateral prefrontal cortex; DMPFC, dorsomedial prefrontal cortex; FOP, frontal operculum; GP, globus pallidus; IFG, inferior frontal gyrus; IPL, inferior parietal lobule; latOFC, lateral orbitofrontal cortex; M1, primary motor cortex; MCC, midcingulate cortex; MDThal, mediodorsal thalamus; MFG, middle frontal gyrus; midMCC, midline midcingulate cortex (no hemispheric assignment); mOFC, medial orbitofrontal cortex; OFC, orbitofrontal cortex; pgACC, pregenual anterior cingulate cortex; PINS, posterior insula; PVG, periventricular gray; S1, primary somatosensory cortex; sgACC, subgenual anterior cingulate cortex; SFG, superior frontal gyrus; SMA, supplementary motor area; SPL, superior parietal lobule; VAThal, ventral anterior thalamus; VLPThal, ventral lateral posterior thalamus; VPLThal, ventral posterolateral thalamus.

While the pain-informative anatomical regions were largely unique within individuals, a subset carried consistent pain-related signals across multiple participants (Fig. 4). Of regions with coverage in at least three of the five above-chance participants (P2 excluded; Supplementary Fig. S1), seven had at least one FDR-significant contact in every covered participant. Most prominently, the right DMPFC replicated across all five of these participants, followed by the left anterior insula, left dorsolateral prefrontal cortex, and left posterior insula (each 4 of 4). Additional regions meeting the criterion (3 of 3) were the right posterior insula and two right-sided thalamic nuclei (mediodorsal (MDThal) and ventral posterolateral (VPLThal)).

We further characterized the directionality of pain-related spectral modulation relative to chronic pain intensity across the top ten features in each participant (Supplementary Fig. S7). In P1, all the top ten features showed increased power during high pain. In contrast, both P3 and P6 showed uniformly decreased power across all top ten features. P4 showed predominantly increased power (8 of 10 features), while P5 exhibited evenly split directionality (5 of 10 increased, 5 of 10 decreased), with the sign of the power change varying across anatomical sites. Together, these results show that pain-related spectral modulations are individualized across participants.

### Pain decoding is enriched in the salience and default mode networks

To better characterize anatomically widespread pain representations, we next asked whether decoding performance is enriched within particular canonical brain networks. We mapped each electrode contact to the Yeo 7-network parcellation using the Schaefer 400-region atlas (see Methods). Across all participants with significant multivariate decoding (P1, P3–P6), 32% of electrode contacts could be assigned to one of these seven networks (using a 7 mm threshold from the nearest region-of-interest (ROI) centroid, robust to 5 mm threshold) (Fig. 5a). Contacts which could not be assigned to a network were excluded from this analysis. Among assigned contacts, the salience network (N = 36), frontoparietal network (N = 17), default mode network (N = 23), and somatomotor network (N = 24) had the most coverage, while the dorsal attention (N = 5) and limbic (N = 1) networks had sparser coverage. No contacts were assigned to the visual network.

**Figure 5.**
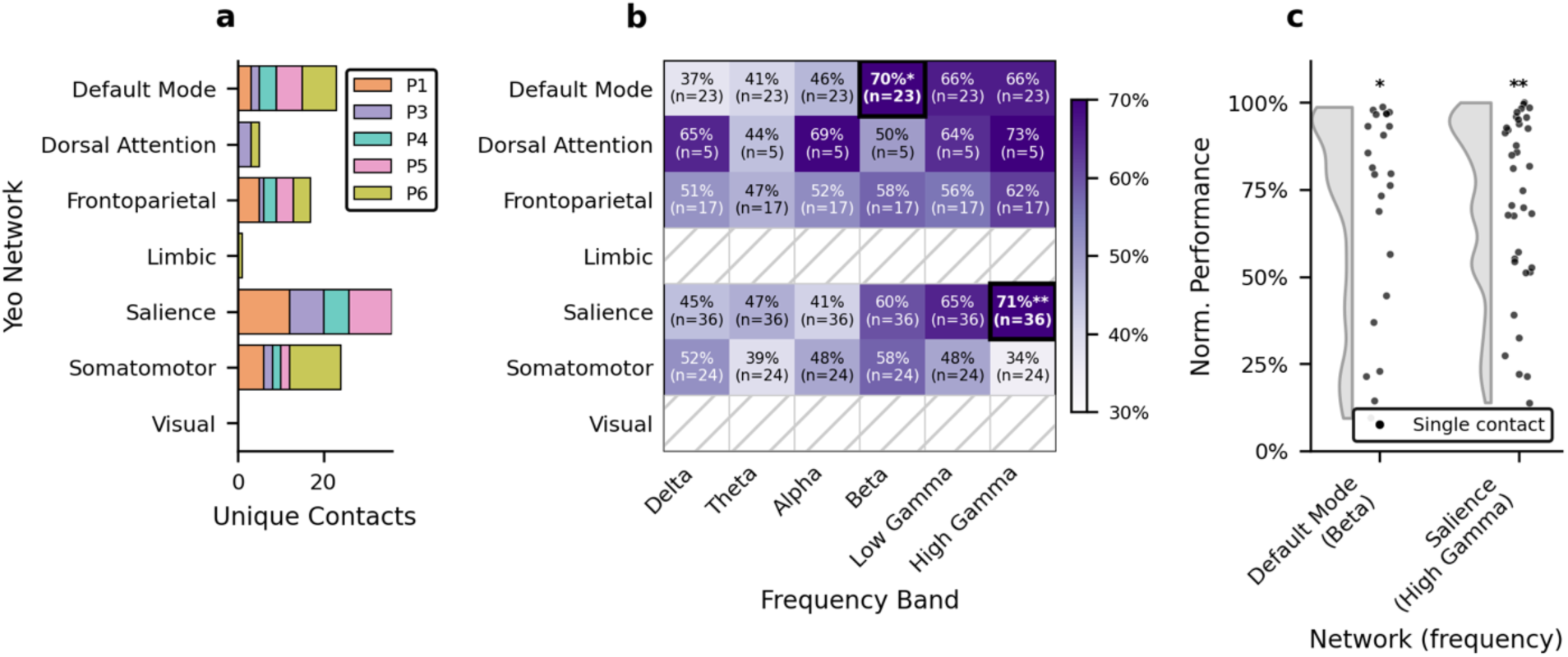
Network-level organization of pain decoding. (a) Breakdown of contacts assigned to each Yeo 7-network parcel, stacked by participant. Limbic and visual were excluded from network-level inference due to insufficient coverage across participants. (b) Heatmap showing the average ranked performance for each network-frequency combination. The average ranked performance and number of contacts within each network are annotated within each cell. Bold cells outlined in black with starred annotations denote network-frequency combinations significant after Benjamini–Hochberg FDR correction across all tested combinations: Default Mode in the beta band and Salience in the high gamma band (*p < 0.05, **p < 0.01). (c) Raincloud plot of normalized per-contact decoding performance within the two FDR-significant network-frequency combinations. Half-violins show the distribution across contacts; dots show individual contacts.

We computed within-participant ranked AUCs normalized from 0 to 1 to prevent higher-performing participants from dominating the pooled network analysis, preserving within-participant contact ordering to ensure that results are not driven by specific participants. We then assessed whether specific network-frequency combinations exceeded chance level using permutation testing that accounted for non-independence of sEEG data within participants (FDR-corrected; see Methods). Two network-frequency combinations reached FDR-corrected significance: (1) salience network in high gamma (mean ranked AUC 0.71, p = 0.006) and (2) default mode network in beta (0.70, p = 0.018) (Fig. 5b, 5c). The enrichment of salience and default mode networks across participants suggests that pain-predictive activity, while locally heterogeneous, may be organized along canonical network boundaries represented in distinct frequency bands.

## Discussion

Using multi-site stereoelectroencephalography recordings in six participants with refractory chronic neuropathic pain, we decoded chronic pain states with a multivariate random forest classifier in five of six participants. Using a leakage-aware Leave-2-Days-Out cross-validation scheme that accounts for temporal autocorrelation, together with control analyses for survey-taking-specific neural activity and long-term drift in neural and pain data, we show that the discovered pain-predictive features track chronic pain states and are unlikely to be fully explained by these confounding variables. We found that pain-predictive signals were distributed across cortical and subcortical foci, with pain-relevant features present in up to 87% of contacts within participants. These results support a distributed, rather than focal, representation of chronic pain in intracranial neural activity. Within this distributed pattern, one region nonetheless stood out: of the regions sampled in every above-chance participant, the right DMPFC was uniquely consistent, carrying significant pain-decoding signal in all five.

Finally, we found strong pain-predictive LFP signatures within the default mode and salience networks in the beta and high-gamma frequency bands, respectively. Prior human neuroimaging studies have identified distributed loci across the brain involved in chronic pain processing, including the anterior cingulate cortex, insula, thalamus, somatosensory cortex, and others. Our univariate analysis supported the inference that pain-predictive signals were broadly distributed across widespread cortical and subcortical regions. The strongest univariate decoders spanned canonical pain-processing regions including the midcingulate cortex (corresponding to dorsal anterior cingulate cortex (dACC) in the pain imaging literature), insula, thalamus, periventricular gray, and prefrontal cortex, with top-performing regions varying across participants. The direction of pain-related spectral modulation was also participant-specific. Despite this variability, the right DMPFC appeared to be a common region harboring biomarker information: every participant whose multivariate decoding was above chance also had at least one FDR-significant pain-decoding contact in this region. The DMPFC’s consistent pain-predictive activity may reflect its well-established involvement in the appraisal and regulation of negative emotion (16,17), processes central to the affective dimension of chronic pain.

Together, these results highlight the need for unique, participant-specific decoders, consistent with a recent study using fMRI-based decoding of widespread chronic pain (10). One possible explanation for interindividual variability is the heterogeneous pain conditions across the cohort, including post-stroke pain (P1, P5), complex regional pain syndrome with degenerative joint disease (P3), spinal cord injury-related pain (P4), chemotherapy-induced neuropathy with degenerative joint disease (P2), and complex regional pain syndrome with post-surgical spine pain (P6). The chance-level decoding performance in P2 may be due to their unique pain etiology. P2’s condition, mixed axial back pain and chemotherapy-induced peripheral neuropathy, may be less associated with central and supraspinal reorganization compared to the other participants’ etiologies. Chemotherapy-induced pain has been conceptualized as a peripheral neuropathic pain condition with prominent peripheral nerve and dorsal root ganglion mechanisms (18), although supraspinal alterations have also been described (19). If P2’s pain is driven predominantly by peripheral mechanisms, it may leave fewer distinctive signatures in our intracranial LFP recordings than pain states with stronger central mechanisms, raising the possibility that decoding from intracranial recordings is most sensitive to centrally mediated pain.

One notable result is the network-level enrichment of pain-decoding performance across the salience and default mode networks, despite the heterogeneity of pain etiologies in our cohort. While individual anatomical labels associated with the strongest univariate decoders varied across participants, the network-level pattern was more consistent. This convergence is consistent with a large resting-state fMRI literature that has reported altered salience and default mode network dynamics across a similarly heterogeneous set of chronic pain diagnoses (20–23) and which has been described to support a triple-network account of chronic pain (7). Intracranial studies have shown that these canonical networks have electrophysiological signatures in human sEEG across multiple independent cognitive tasks (24). Our results extend this framework to chronic pain, suggesting that the network-level signatures identified by fMRI have a decodable mesoscale electrophysiological complement in human intracranial local field potential signals. That pain-informative signals were prominent in the gamma band suggests that this activity reflects summed extracellular postsynaptic potentials from co-firing of neurons distributed across millimeters of cortex (25). This mesoscale origin makes it a particularly appealing marker for processes involving integration of activity across multiple brain regions. Within the present chronic pain cohort, the salience and default mode networks showed distinct spectral profiles, with salience network signals peaking in the high-gamma band and default mode network contacts peaking in the beta band. Beta-band activity has been proposed to carry top-down feedback signals along the cortical hierarchy that actively maintain ongoing cognitive and sensorimotor states (26–28). A beta signature localized to default mode network nodes would therefore be consistent with the idea that the chronic pain state is actively sustained by top-down processes within high-order association cortex, rather than by ongoing bottom-up nociceptive drive.

Identifying pain-relevant signals within cortical networks has direct translational implications. Current DBS approaches for chronic pain rely on deep brain targets such as the thalamus and periventricular gray, as well as cortical targets such as the anterior cingulate cortex and insula, which require invasive stereotactic implantation. The enrichment of pain-related signals within cortical networks suggests that superficial cortical targets accessible via electrocorticography as well as less invasive approaches, such as transcranial magnetic stimulation or focused ultrasound, may also facilitate closed-loop therapies. For depth-based approaches, emerging electrode designs with more contacts per lead, along with multi-lead trajectories, may allow cortical pain-decoding sites to be co-specified with subcortical stimulation targets during trajectory planning, enabling both to be reached with a single implant. However, these network-level results should be interpreted with caution. Only 32% of gray matter contacts could be assigned to a canonical Yeo network, and several networks, including the limbic, dorsal attention, and visual networks, had too few electrodes for meaningful testing. Because per-participant coverage within any single network was sparse, our network analyses pool contacts across participants and do not capture within-participant functional connectivity. Future work with denser within-network coverage should examine whether connectivity metrics, such as coherence or phase synchrony, carry additional pain-state information.

This study has limitations. Our sample of six participants with mixed pain etiologies limits the generalizability of our findings. Pain scores were binarized to reflect the threshold-crossing paradigm used in closed-loop DBS devices, but this discards information about graded pain intensity that continuous decoding approaches could capture. More broadly, we did not control for other potential confounds such as arousal or medication timing, both of which could covary with pain over multi-day recordings. Our Leave-2-Days-Out cross-validation and detrending analyses already mitigate within-day medication cycles and slow drifts, while the cross-context analysis argues against confounds tied to survey-taking arousal. Finer-grained controls for arousal state and medication regimen will be valuable in future work. Additionally, coverage was not brain-wide, and subcortical structures could not be mapped onto the Yeo cortical parcellation, leaving the network-level organization of subcortical pain signals uncharacterized. Future work with larger cohorts, matched pain etiologies, and denser electrode coverage will be needed to determine whether the network-level convergence observed here generalizes across individuals and pain conditions.

In conclusion, using a leakage-aware cross-validation approach, we demonstrate that spontaneous chronic pain states are decodable from widespread intracranial neural activity over multi-day timescales. Pain-predictive features were broadly distributed across cortical and subcortical structures and were individualized across participants but were preferentially enriched within the salience and default mode networks at the macroscale. These findings provide the first network-level characterization of how oscillatory neural activity represents clinically relevant chronic pain in intracranial recordings, and offer a foundation for informing personalized, multi-site closed-loop neuromodulation therapies.

## Methods

### Participants and clinical protocol

Six participants with chronic refractory neuropathic pain were included in the study (P1–P6, aged 48–58, 3 female). Participants were enrolled in a clinical trial for personalized closed-loop deep brain stimulation (ClinicalTrials.gov: NCT04144972) (3); the present analysis uses neural data from the temporary inpatient sEEG mapping phase. P6 is an additional participant enrolled in the same trial but not included in the prior report. Participants had heterogeneous chronic neuropathic pain etiologies: P1 and P5 had chronic post-stroke pain; P2 had chemotherapy-induced neuropathy and degenerative joint disease of the cervical and lumbar spine; P3 had complex regional pain syndrome and degenerative joint disease with radiculopathies in the cervical and lumbar spine; P4 had spinal cord injury-related pain; and P6 had complex regional pain syndrome and spine pain resulting from complications of a prior spinal cord stimulator implantation surgery. All participants provided written informed consent.

During the trial, participants underwent an 8–10 day monitoring period during which stereoelectroencephalography (sEEG) electrodes were implanted. During this time, neural data were continuously recorded, brain stimulation parameters were tested, and pain surveys were collected. Pain intensity was recorded using a visual analog scale on a tablet (VAS, 0–100) multiple times daily.

### Electrode implantation and localization

Stereoelectroencephalography electrodes were implanted bilaterally, targeting cortical and subcortical structures implicated in pain processing. Targeted regions included the anterior cingulate cortex (ACC), midcingulate cortex (MCC), insula, dorsolateral prefrontal cortex (DLPFC), orbitofrontal cortex (OFC), primary somatosensory cortex (S1), primary motor cortex (M1), supplementary motor area (SMA), ventral posterolateral thalamus (VPLThal), centromedian thalamus (CMThal), periaqueductal/periventricular gray (PAG/PVG), caudate, and globus pallidus. In addition, four participants (P1–P4) had electrocorticography (ECoG) strips placed over primary motor and somatosensory cortices.

Electrode localization was performed manually. First, CT and MRI images were co-registered. Following this, contact labels were manually assigned. Only gray matter contacts were included in all subsequent analyses, yielding 57–90 contacts per participant. Regions labeled as midcingulate cortex (MCC) in our anatomical parcellation correspond to what is often referred to as dorsal anterior cingulate cortex (dACC) in much of the pain neuroimaging literature (29,30).

### Pain assessment and binarization

Participants reported pain intensity on a visual analog scale from 0 to 100, with the question being: “Please rate your current pain intensity.” For classification analysis, continuous pain scores were binarized within each participant, with the top 33% of pain scores being considered high pain and the bottom 33% low pain. Scores falling between these thresholds were excluded from analysis. Binary classification was used to reflect the threshold-crossing paradigm used in current closed-loop DBS devices. The decision to use a 33rd percentile threshold was made to ensure an equal balance of high and low pain scores, while maintaining a consistent approach across participants.

### Neural data acquisition and preprocessing

Signals were acquired using a Nihon Kohden 256-channel amplifier with a single reference contact located in a confirmed white matter tract. Data were sampled at either 2 or 10 kHz. Depth electrodes (PMT or ADTech), each with 8–14 cylindrical contacts spaced 5 mm apart, were used for sEEG recording. Subdural ECoG strips containing 8–20 contacts were used in four participants (P1–P4).

Signals were first re-referenced using a Laplacian montage, in which each contact’s signal was replaced by the difference between its potential and the average of its immediate neighbors on the same electrode shaft. For contacts at the ends of an electrode, the single adjacent contact was used. This approach attenuates volume-conducted activity common to nearby contacts and enhances the spatial specificity of the recorded signals. Line noise and its harmonics were removed using a notch filter at 60, 120, 180, and 240 Hz. A bandpass filter (0.1-250 Hz, finite impulse response) was then applied to remove slow drift and high-frequency noise. Finally, signals were downsampled to a common sampling rate of 1,000 Hz to standardize the temporal resolution across participants and reduce computational load.

sEEG signals were continuously recorded throughout the inpatient trial. For each of the pain surveys collected, 30 seconds of neural data were extracted after the survey start. This window was chosen to approximate the minimum survey completion time observed across participants. We chose to analyze during-survey data as participant behavior was consistent between high and low pain states (they were always taking a survey), whereas pre-survey behavior was less controlled. Additionally, a 30-second epoch preceding the survey was extracted to ensure that the pain signals generalized beyond survey-taking behavior (−60 to −30 seconds relative to survey onset). Data recorded within 5 minutes after stimulation were excluded from this analysis. Five minutes was used as a data-driven compromise between maximizing the number of surveys to be used in the analysis and avoiding contamination from stimulation.

### Artifact rejection and epoch selection

Recording artifacts in the during-survey sEEG epochs were identified using an automated pipeline as described in (31). Each of the 30-second epochs was divided into six non-overlapping 5-second windows. Each window was then flagged as noisy or clean by the classifier. Trials with fewer than four remaining clean windows were discarded. From the surviving clean windows, the last four, corresponding to the latter 20 seconds of the epoch, were retained.

### Spectral feature extraction

Spectral power features were extracted using a multitaper fast Fourier transform (time-bandwidth product TW = 2, L = 3 tapers). After artifact rejection, the mean spectral power across the four clean windows was calculated in each of six canonical frequency bands: delta (0.5–4 Hz), theta (4–8 Hz), alpha (8–12 Hz), beta (12–30 Hz), low gamma (30–70 Hz), and high gamma (70–150 Hz).

### Multivariate decoding pipeline

A multivariate random forest classifier was used to decode binary pain states from the neural data. Specifically, the features were region-level spectral power as extracted using the PCA approach outlined in Figure 2a and below. Leave-2-Days-Out cross-validation was used.

#### Cross-validation scheme

Both chronic pain and neural signals are known to exhibit temporal autocorrelation (2,32). Samples close in time are not independent and share variance beyond what is captured by the pain label alone. Standard cross-validation such as random k-fold or leave-one-out can therefore lead to train-test leakage by, for example, allowing a classifier to memorize features of surveys on either side of a held-out sample. To guard against this, we employed a Leave-2-Days-Out cross-validation scheme (Supplementary Fig. S3). For every possible combination of two recording days, we held out those two days as the test set and trained using the remaining days. This ensured that training and testing data never occurred close together in time. The two-day holdout approximated a roughly 80/20 train-test split, in line with standard practice. Only folds in which both pain classes were present in the held-out days were retained for evaluation; folds containing only a single class were excluded, as AUC is undefined in that case. The final per-participant performance was computed as the mean AUC across all retained folds.

#### Feature construction

We employed feature reduction to avoid model overfitting. Features were constructed separately for each Leave-2-Days-Out cross-validation fold. Spectral power features were first grouped by anatomical region and frequency band. For each anatomical region, features were z-scored using the training set mean and standard deviation. Principal component analysis (fit on training data) was then applied across all contacts within each frequency band and anatomical region, retaining the first principal component. Reducing the signals from a single anatomical region to one feature served two purposes. First, it ensured an equal number of features per anatomical target regardless of contact count, and it also reduced feature dimensionality to mitigate the potential for overfitting given the limited number of trials. All features were then concatenated and rescaled using training data to form the final feature vector.

#### Classification and evaluation

A random forest classifier (500 trees, balanced class weights) was trained on the dimensionality-reduced features. Performance was then evaluated using the area under the receiver operating characteristic curve (AUC), which was calculated for each of the Leave-2-Days-Out folds. The primary metric used to report performance is the mean AUC across all folds. To visualize per-survey performance, and because individual samples were present across multiple cross-validation folds, predictions were aggregated across folds before computing the performance. Raw predicted probabilities can vary in scale across folds due to differences in training set composition. To address this issue, we first rank-transformed the predicted probabilities within each fold and rescaled them to [0,1] by dividing by the fold size. Next, for each sample, the ranked probability was averaged across folds. The optimal classification threshold was then determined using Youden’s J statistic, which maximizes the sum of sensitivity and specificity on the ROC curve of the ranked predictions (33). Samples with an averaged ranked probability at or above this threshold were classified as high pain. Permutation feature importance was used to evaluate neural contributions to model performance. For each Leave-2-Days-Out fold, each post-PCA region-frequency feature was randomly shuffled 30 times on the held-out test set, and the decrease in AUC relative to the unshuffled prediction was recorded. Per-feature importance was then averaged across folds.

#### Permutation testing

To test for statistical significance, permutation testing was employed in which pain labels were shuffled 1,000 times and the full pipeline (feature scaling, PCA, classification) was recalculated. We define the permutation p-value as the proportion of null AUCs exceeding the observed AUC. All p-values were corrected for multiple comparisons across participants using the Benjamini-Hochberg false discovery rate (FDR) procedure.

#### Control analyses

Because both pain reports and neural features may drift over multi-day inpatient recordings, we repeated the decoding analysis after removing linear temporal trends from the spectral features. The Leave-2-Days-Out cross-validation scheme alone does not account for multi-day trends as it only ensures that within-day samples never co-occur in the training and testing sets, whereas these long-term trends cause leakage between neighboring days. The same permutation procedure outlined above was used to test for detrended model significance. To test whether pain signatures generalized beyond survey-taking behavior (such as introspection), we performed the decoding analysis by training on during-survey data but testing on pre-survey data.

#### Multivariate feature multicollinearity

Significant correlation among features would impede the meaningful interpretation of feature importances, as the contribution of any single feature cannot be isolated when correlated features are removed. To test for multicollinearity, the region-level PCA approach was applied across the whole dataset in the same fashion as the decoding pipeline. We then calculated the Pearson correlations among the features and characterized the distribution of absolute correlation coefficients. Furthermore, effective dimensionality analysis was employed in which we determined the number of principal components needed to explain 95% of the total variance of the features, which we expressed as a percentage of the total feature count. A lower effective dimensionality relative to the total feature count would indicate substantial redundancy among the features, whereas a higher percentage would suggest that the PCA-reduced features capture largely independent sources of variance.

#### Univariate contact-level decoding

To interpret the spatial organization of electrode contacts containing pain-relevant information, we employed contact-level univariate logistic regressions. Each gray matter contact and frequency band combination was independently used to predict chronic pain intensity using Leave-2-Days-Out cross-validation. Within each fold, power was z-scored using the training set mean and standard deviation and used to train an L2-regularized logistic regression classifier. AUC values were collected within each fold and then averaged to yield the final evaluation metric.

To test for significance, pain labels were shuffled 1,000 times per participant and the full pipeline was re-run on every contact-frequency feature. For each feature, a null AUC distribution was constructed from the resulting 1,000 permutations, and a feature-specific significance threshold was defined as the 95th percentile of that distribution. Per-contact significance was assessed by Benjamini–Hochberg FDR correction across the six frequency-band p-values, with a contact labeled significant if at least one band survived correction. Multiple comparisons correction was applied across frequency bands within each contact, not across all contacts within a participant. For visualization of participant-level frequency composition, we computed the proportion of each participant’s FDR-significant features falling in each band (within-participant normalization).

#### Directionality analysis

To describe the sign of pain-related spectral changes among the strongest local decoders, we selected each participant’s top 10 contact-frequency univariate features ranked by cross-validated AUC. For each selected feature, directionality was defined by the difference in mean spectral power between high-pain and low-pain observations, such that positive values indicate greater power during high pain and negative values indicate lower power during high pain. This analysis was intended as a descriptive summary of the top-performing features rather than a separate inferential test.

#### Network-level analysis

To test whether pain-decoding contacts were organized within canonical cortical networks, gray matter contacts were first mapped to the Yeo 7-network parcellation using the Schaefer 400-region cortical atlas (15,34). The nearest Schaefer ROI for each contact was identified in Montreal Neurological Institute (MNI) space using Euclidean distance. Contacts were considered part of a cortical network if located within 7 mm of the nearest ROI centroid; electrode contacts not assigned to a network were excluded from this analysis. This threshold was chosen to account for co-registration error and the spatial extent of sEEG contacts; results were robust to a more conservative 5 mm threshold. P2 was excluded from network-level analyses due to non-significant decoding performance. To prevent higher-performing participants from dominating the pooled network analysis, within-participant contact AUCs were rank-transformed and rescaled to [0,1], preserving within-participant ordering while removing between-participant differences in overall decoder strength.

#### Network-level permutation testing

To assess whether specific network-frequency combinations exhibited above-chance decoding, we employed a permutation test that accounted for the non-independence of sEEG data recorded from the same participant. Ranked AUC values were first shuffled within each participant across 10,000 permutations, preserving the participant-level correlation structure while breaking the association between AUC values and network assignments. Within each permutation, the mean ranked AUC was recomputed for all network-frequency combinations. The p-value was defined as the proportion of 10,000 permutations in which the permuted mean ranked AUC equaled or exceeded the observed mean ranked AUC. Only network-frequency combinations with at least five electrode contacts were tested. P-values were corrected across all tested network-frequency combinations using the Benjamini-Hochberg FDR procedure (alpha = 0.05).

## Supplementary Information

**Supplementary Figure S1.**
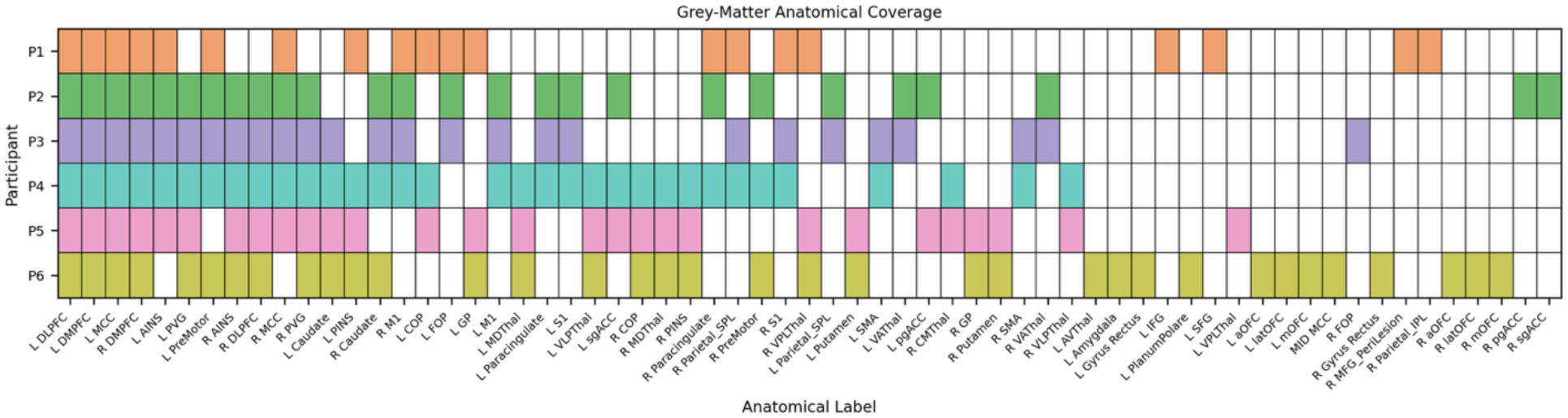
Gray-matter anatomical coverage per participant. Binary coverage map indicating whether each participant (rows, P1–P6) had at least one gray-matter sEEG contact in each anatomical region (columns). Filled cells (colored by participant) indicate coverage; empty cells indicate no coverage. Columns are sorted in descending order of coverage prevalence across the cohort.

**Supplementary Figure S2.**
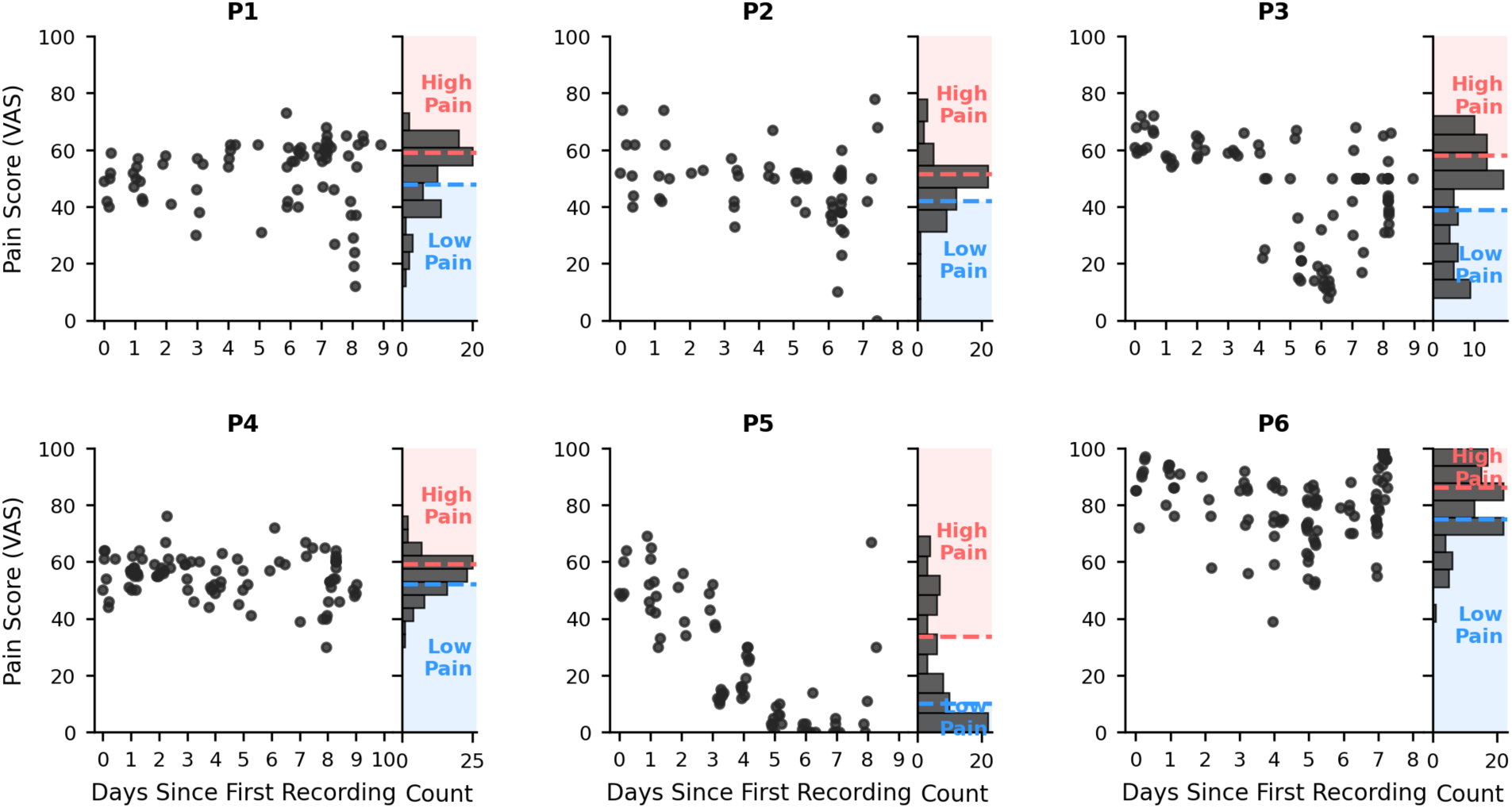
Per-participant pain ratings and classification thresholds. Scatter plots (left sub-panels) show the VAS pain ratings over the recording period for each participant (P1–P6). Histograms in the margins show the counts for these pain scores, as well as the thresholds used to define high (red) vs. low (blue) pain for classification (top/bottom 33rd percentile, respectively). The middle 33% of scores were excluded from decoding analysis.

**Supplementary Figure S3.**
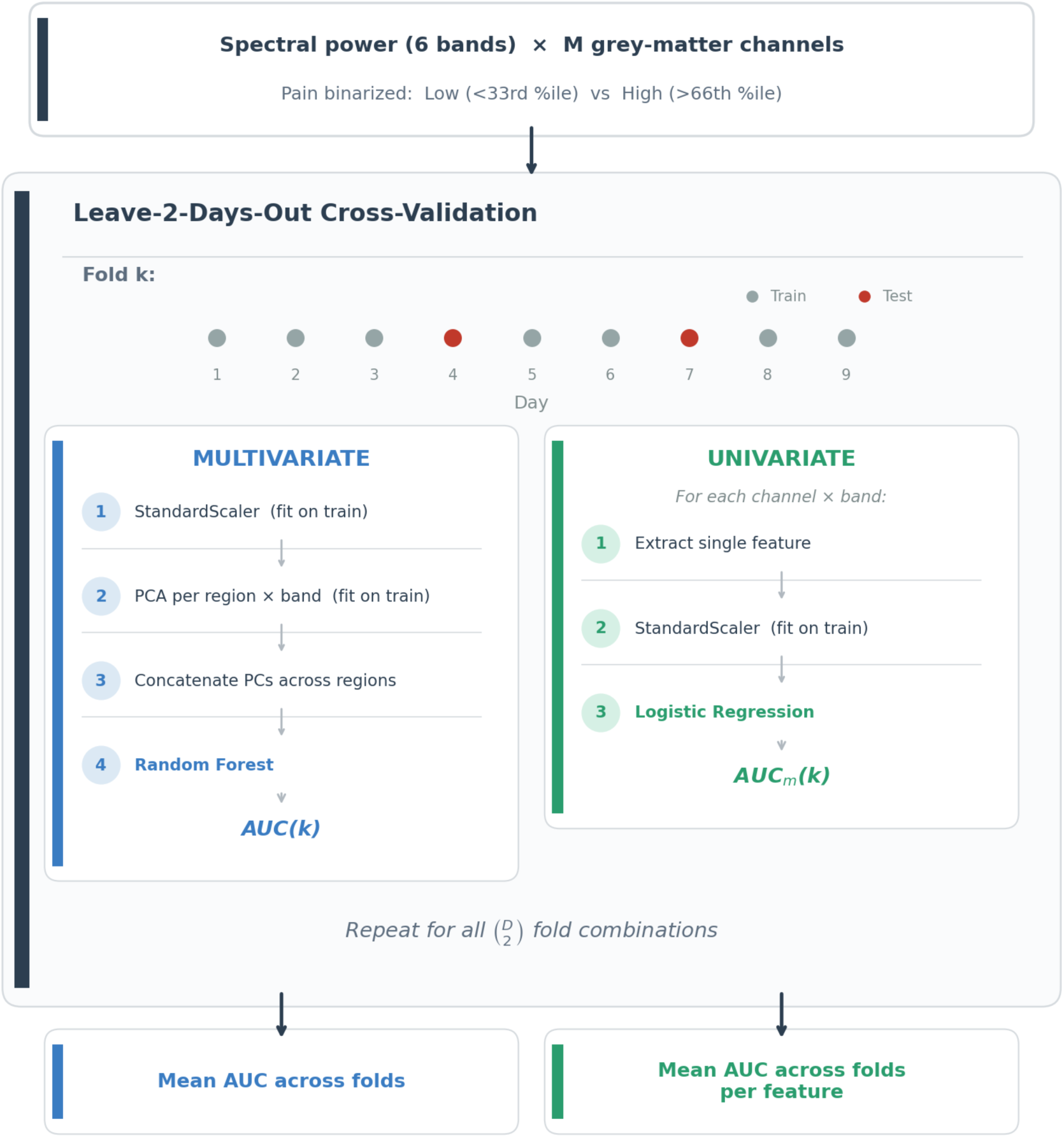
Cross-validation and decoding pipeline overview. A Leave-2-Days-Out cross-validation scheme was designed to account for temporal autocorrelation in the neural data and pain scores, which can lead to train-test leakage. Per-contact spectral power features from six frequency bands and VAS pain labels are used. For all possible combinations of two days (example fold shown in figure), multivariate and univariate approaches were independently employed. Multivariate (left): feature reduction was used in the multivariate classifier to prevent overfitting due to a large number of features compared to the number of samples; ultimately, this method yields one feature per anatomical region and frequency band. Univariate (right): for each contact-frequency combination, a separate univariate logistic regression classifier was trained to test for the predictive power of single contacts. After decoding performance was calculated for all (D choose 2) folds, where D = the number of days, results were averaged across folds to yield the final per-participant AUC.

**Supplementary Figure S4.**
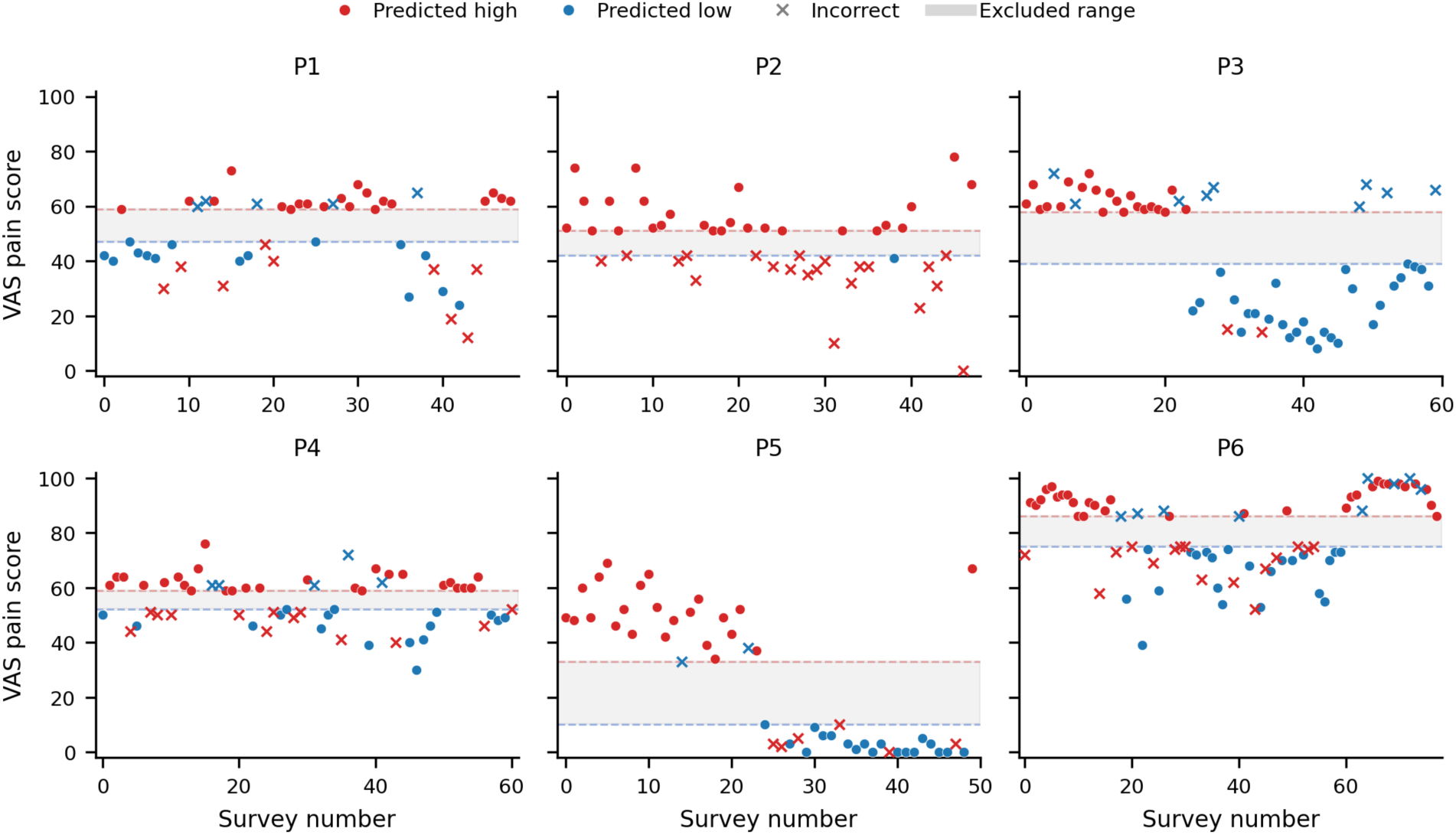
Per-participant multivariate decoder predictions. Each panel shows all VAS pain scores for one participant (P1–P6). Surveys are colored by the multivariate random forest decoder’s predicted class: red for predicted high pain, blue for predicted low pain. Misclassified surveys are marked with an x. Dashed lines indicate the participant-specific classification thresholds (blue, 33rd percentile; red, 67th percentile), and the gray shaded band marks the excluded middle range of scores not used in decoding.

**Supplementary Figure S5.**
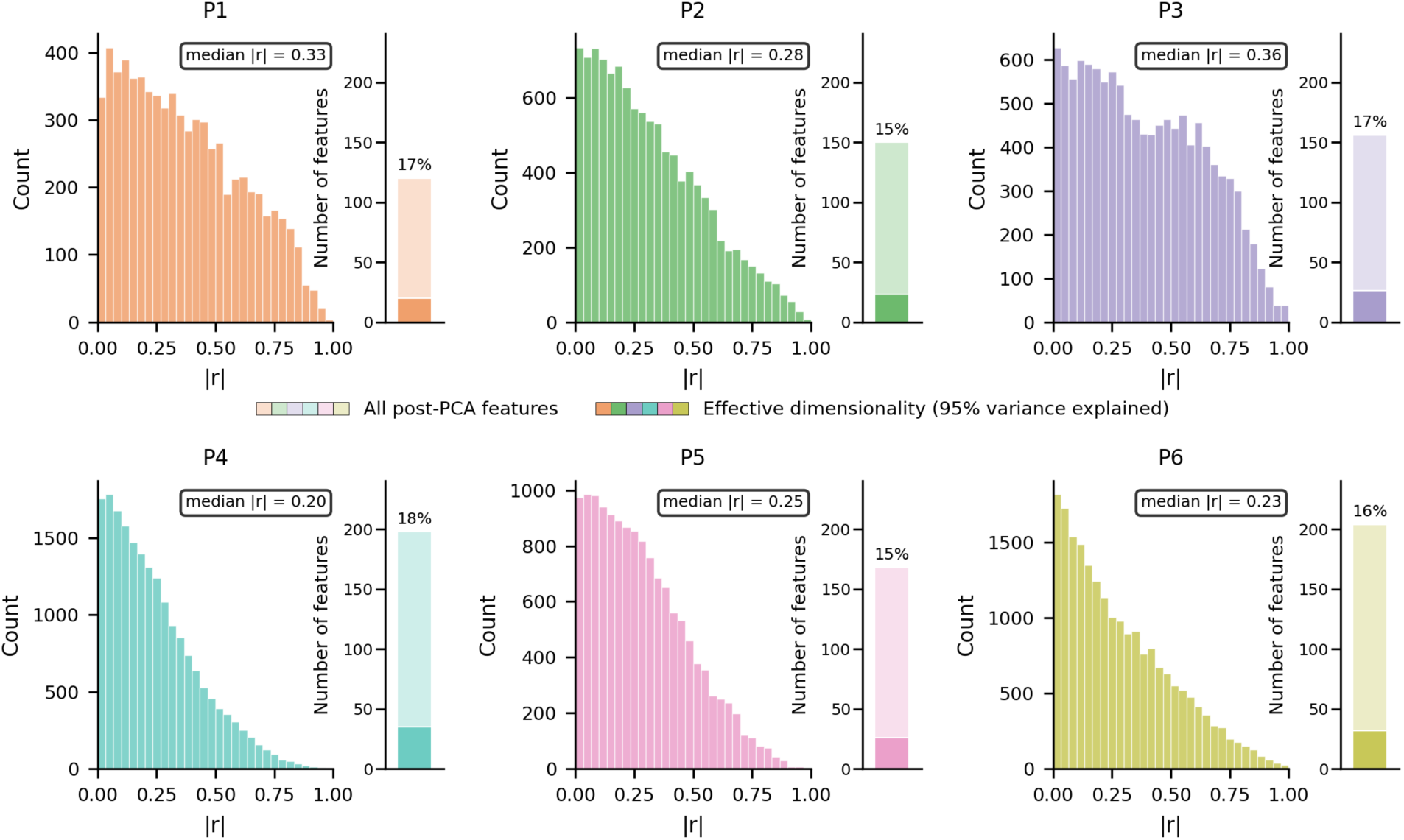
Feature correlation structure and effective dimensionality. Per-participant histograms of absolute pairwise Pearson correlations (|r|) computed across all post-PCA features used in the multivariate classifier (P1–P6). Median |r| is shown in each panel. Inset stacked bar: total number of post-PCA features (light shading), and the effective dimensionality, defined as the number of principal components needed to explain 95% of the variance (dark shading); percentage label shows the ratio of effective to total dimensionality. Across participants, effective dimensionality ranged from 15% to 18% of the post-PCA feature count, reflecting substantial feature redundancy.

**Supplementary Figure S6.**
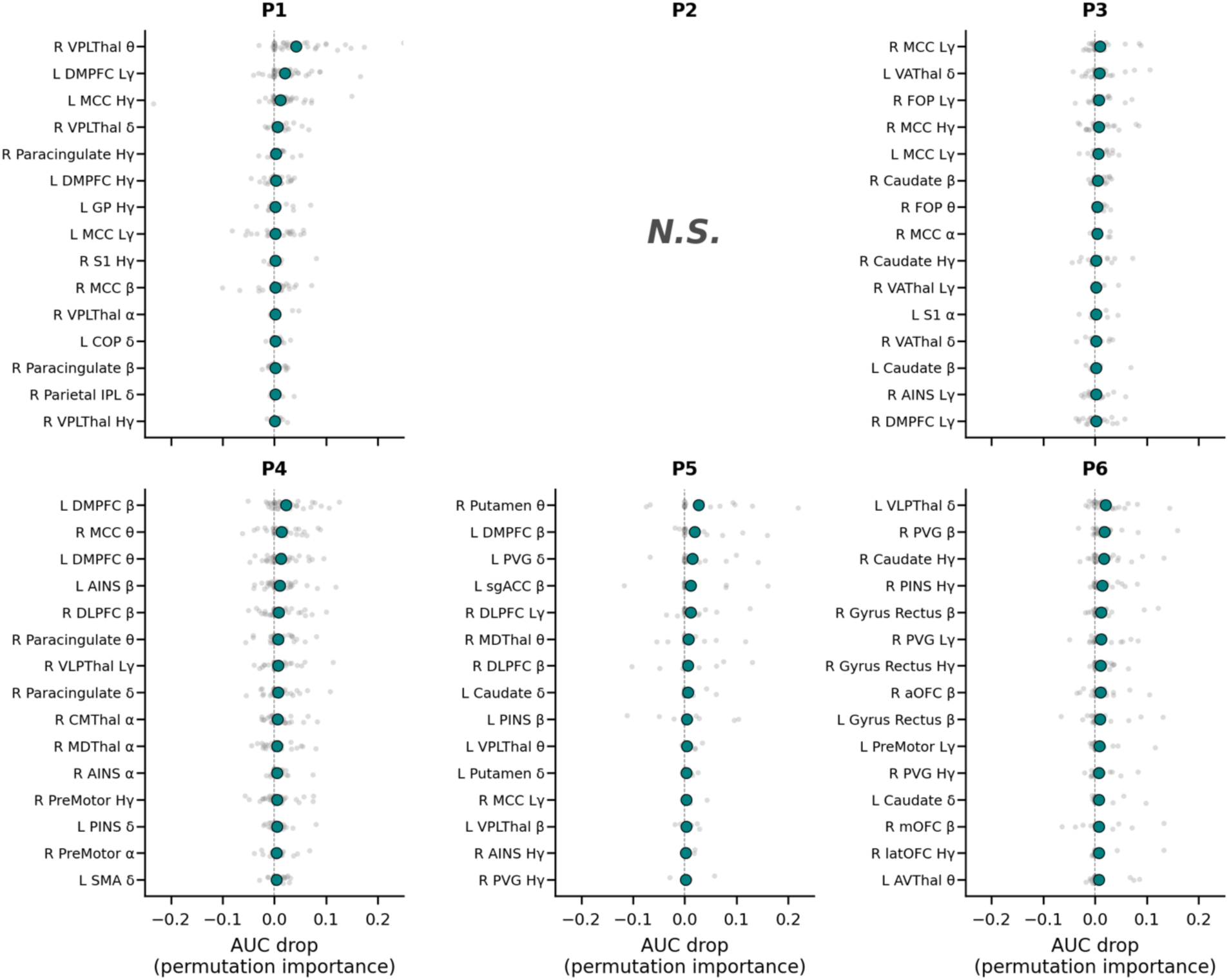
Permutation feature importance for the multivariate decoders. Top 15 features for each participant ranked by permutation importance (change in AUC when shuffling that individual feature). Teal points, mean across cross-validation folds; gray points, individual fold estimates. Each feature is the result of the within-region-frequency PCA reduction described in Methods. Small AUC drops indicate feature redundancy: informative content can be substituted by correlated features in the model (see Supplementary Fig. S5), so low importance does not necessarily mean a feature is uninformative. P2 is shown as N.S. because decoding did not reach significance.

**Supplementary Figure S7.**
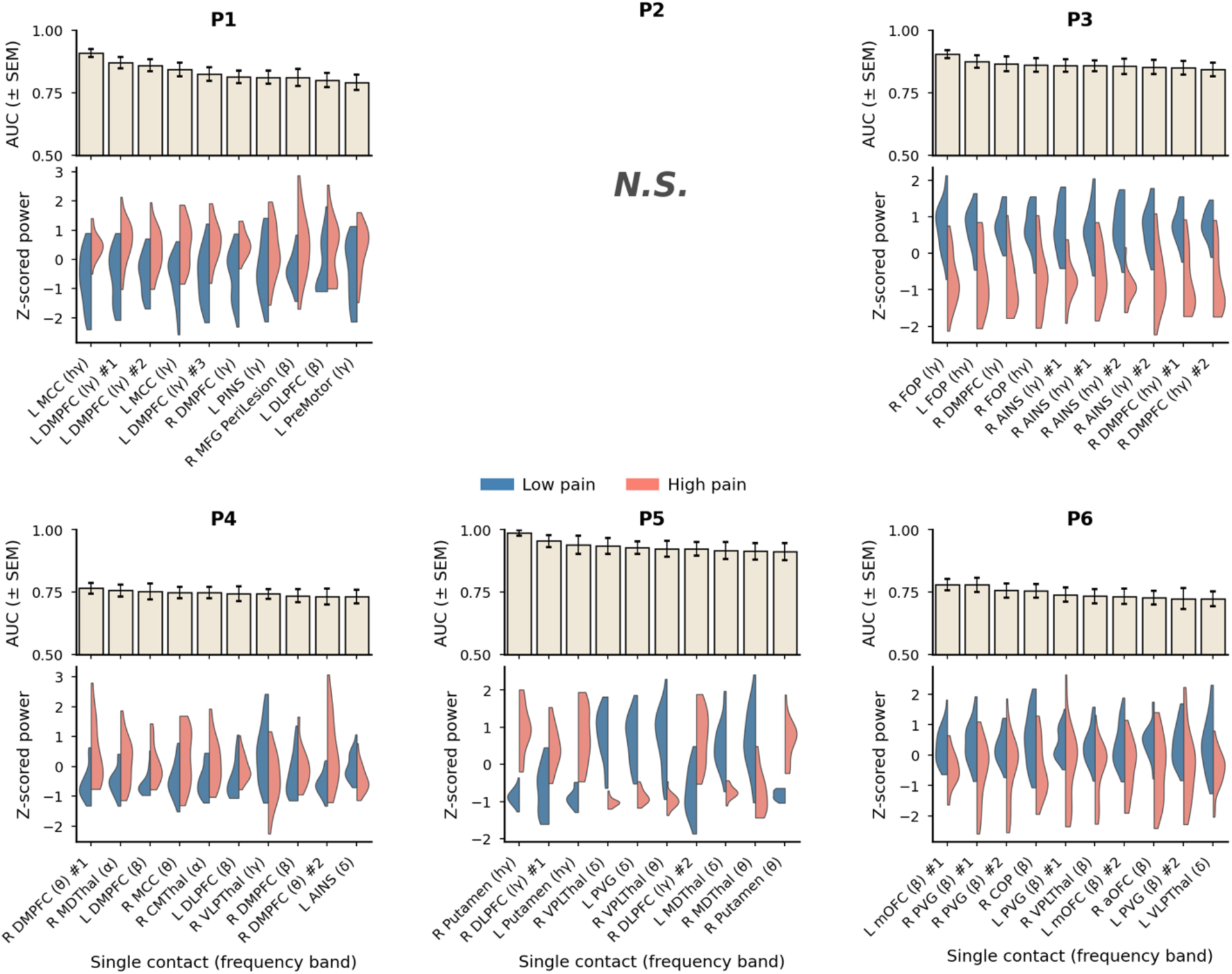
Top-performing single-contact univariate decoders. Per-participant summary of top 10 contact-frequency univariate classifiers. Top panels: mean AUC ± SEM across cross-validation folds for each single-feature classifier. Bottom: Split-violin showing the distribution of feature power values for high (red) vs. low (blue) pain states. Feature labels include the anatomical region each contact is within. It is possible that multiple contacts can have strong predictive features in the same frequency band and anatomical region; in this case, a number has been placed after the feature label to denote individual contacts. P2 did not reach significance so no features are shown. P5 results should be interpreted considering the temporal-confound caveat described in the main text.

**Supplementary Table S1.** Participant characteristics and pain survey statistics. For each participant (P1–P6): Number of surveys, total number of completed pain surveys; Days, length of the inpatient recording period in days; Median VAS [IQR], median pain intensity on the visual analog scale (0–100) with interquartile range; Low/High Pain VAS Threshold, participant-specific 33rd / 67th VAS percentiles used for binary classification; Number of High/Low Pain Samples, number of surveys falling below the 33rd percentile (low pain) and above the 67th percentile (high pain), respectively.

| Participant | Number of surveys | Days | Median VAS [IQR] | Low/High Pain VAS Threshold | Number of High Pain Samples | Number of Low Pain Samples |
| --- | --- | --- | --- | --- | --- | --- |
| P1 | 72 | 10 | 55.5 [42.8–60.2] | 47.0 / 59.0 | 25 | 26 |
| P2 | 57 | 8 | 50.0 [40.0–52.0] | 42.0 / 51.0 | 26 | 22 |
| P3 | 88 | 10 | 50.0 [31.0–60.0] | 39.0 / 58.0 | 30 | 30 |
| P4 | 89 | 10 | 56.0 [50.0–60.0] | 52.0 / 59.0 | 31 | 30 |
| P5 | 72 | 9 | 15.0 [4.5–43.0] | 10.0 / 33.0 | 25 | 25 |
| P6 | 105 | 8 | 82.0 [73.0–90.0] | 75.0 / 86.0 | 40 | 38 |

